# Development of vascular myogenic responses in zebrafish

**DOI:** 10.1101/713248

**Authors:** Nabila Bahrami, Sarah J. Childs

**Affiliations:** Alberta Children’s Hospital Research Institute and Department of Biochemistry and Molecular Biology, University of Calgary, 3330 Hospital Drive NW, Calgary, Alberta, Canada, T2N 4N1

**Keywords:** pericytes, vascular smooth muscle cells (vSMCs), blood vessel, myogenic tone, Zebrafish

## Abstract

The vascular system is placed under enormous stress at the onset of cardiac contractility and blood flow. Nascent blood vessel tubes initially consist of a thin endothelial wall and rapidly acquire support from mural cells (pericytes and vascular smooth muscle cells; vSMCs). Following their association with vessels, mural cells acquire vasoactive ability (contraction and relaxation). However, we have little information as to when this vasoactivity first develops, and the extent to which each mural cell type contributes to vascular tone regulation during development. For the first time in an in vivo system, we highlight the dynamic changes in mural cell vasoactivity during development. We assess mural cell vasoactivity in the early zebrafish cerebral vasculature in response to pharmacological agents. We determine that pericyte-covered vessels constrict and dilate at 4 days post fertilization (dpf) but not at 6 dpf. The prostaglandin EP4 receptor contributes to pericyte-covered vessel dilation at 4 dpf. In contrast, vSMC-covered vessels constrict but do not dilate at 4 dpf. At 6 dpf, vSMC-covered vessels continue to constrict but only dilate from a pre-constricted state. Using genetic ablation, we demonstrate that mural cell contraction and relaxation is an active response by pericytes and vSMCs. Thus, we show that both pericytes and vSMCs have the ability to regulate cerebral vascular tone but at different stages of development. Pericytes are involved in regulating vessel diameters prior to the maturation of the vSMCs. Once vSMCs mature, pericytes are no longer active, and only vSMCs regulate vascular tone in the developing embryonic brain of zebrafish. The onset of vasoactivity of vSMCs corresponds to the development of increased neuronal activity and neurovascular coupling.

## INTRODUCTION

The cerebral vascular network is stabilized by perivascular cells, which include pericytes, vascular smooth muscle cells (vSMCs), astrocytes, microglia, perivascular macrophages (PVMs) and fluorescent granular perithelial cells (FGPs; originally known as Mato cells (Gaengel et al., 2009; Galanternik et al., 2017; Mato and Ookawara, 1979; Williams et al., 2001).Two of these cell types, pericytes and vSMCs, have additional roles in regulating vessel diameter through active contraction and relaxation.

We are using zebrafish to study the function of mural cells in vascular regulation during development. Pericytes and vSMCs of the head and trunk in zebrafish are of different developmental origins. Consequently, there may be differences between anatomical locations and mural cell function. Pericytes of the trunk come from the sclerotome (Ando et al., 2016; Arciniegas et al., 2000; Santoro et al., 2009; Stratman et al., 2017) while those of the head originate from both the neural crest and the mesoderm (Ando et al., 2016; Cavanaugh et al., 2015; Kok et al., 2015; Wang et al., 2014; Whitesell et al., 2014). We have insights into vascular tone regulation in the zebrafish trunk. The dorsal aorta (DA) acquires tone between 48 and 80 hours post fertilization (hpf) and the diameter of the DA varies inversely with vSMC number (Stratman et al., 2017). Furthermore, platelet-derived growth factor receptor beta (*pdgfrβ*) regulates the development of vascular tone in the zebrafish trunk (Stratman et al., 2017), likely through regulating mural cell differentiation. However, there is no information about the acquisition of vasoactivity in brain vessels during development.

The process of mural cell recruitment to the endothelium is still poorly understood and there are conflicting reports. In zebrafish, when blood flow initiates at approximately 26 hpf, it induces shear stress on the arterial endothelium. In this low-flow state, shear stress causes deflection of endothelial cilia that function as mechanosensors, and induce the recruitment of mural cells to the endothelium (Chen et al., 2017; Goetz et al., 2014). However, an independent study found that mural cell recruitment is flow-independent (Ando et al., 2016) around the basilar artery (BA) and central arteries (CtAs) of the brain. Also, a lack of blood flow did not affect mural cell development on the dorsal aorta (DA) and intersegmental vessels (ISVs). Our understanding of mural cell recruitment is still evolving.

Once attached to the endothelium, pericytes are generally solitary cells. Their morphology varies along the vascular tree but they all have a rounded cell body and long processes, and attach to the longitudinal axis of capillaries (Armulik et al., 2005; Armulik et al., 2011; Attwell et al., 2016; Gaengel et al., 2009; Hartmann et al., 2015). Pericytes share the basement membrane with endothelial cells and the two cell types physically contact each another via peg and socket connections (Armulik et al., 2005; Armulik et al., 2011). In contrast, vSMCs wrap perpendicularly around vessels. They are connected to endothelial cells via myoendothelial gap junctions, allowing for direct signals from one cell to the other (Borysova et al., 2018). Vessel size determines the degree of coverage by vSMCs as they form multi-layers around larger vessels and a single layer around smaller vessels (Santoro et al., 2009).

*pdgfrβ* marks pericytes in zebrafish and is one of the earliest pericyte markers (Ando et al., 2016; Wang et al., 2014). *pdgfrβ* transgene expression can be seen as early as 8-somite stage in the neural crest and marks cerebral pericytes by 48 hpf. Cerebral vSMCs can be visualized using the transgenic line *Tg*(*acta2:EGFP*, (Whitesell et al., 2014), which is expressed in smooth muscle cells and is first seen at about 3-3.5 days post fertilization (dpf) in the ventral head, much later than the pericyte marker.

In vitro, in vivo, and ex vivo approaches in the adult mouse model have shown that while there is robust control of vascular diameter by vSMCs, pericytes have more limited and controversial roles. The contraction of pericytes was first largely based on indirect evidence (Díaz-Flores et al., 2009; Hamilton et al., 2010; Puro, 2007). For instance, pericytes were found to contract during ischemia and remain contracted even after reperfusion of the occluded artery (Yemisci et al., 2009). Other studies found that although pericytes are contractile and capable of modulating cerebral blood flow (CBF), they do not play a major role in the process of neurovascular coupling (Fernandez-Klett et al., 2010). However, in some of these early studies, pericytes were identified by their morphology without the use of markers. Using adult rat brain slices and in vivo methods in the mouse brain, Hall et al. found that pericytes rather than vSMCs regulate cerebral vessel diameters and the consequent changes in red blood cell (RBC) velocity (Hall et al., 2014). They found pericyte-covered capillaries to dilate prior to vSMC-covered arterioles. However, others have shown that smooth muscle actin is only present in mural cells with a smooth muscle morphology and not in those with a pericyte morphology (Hill et al., 2015). Using the adult mouse cerebral cortex, Hill et al. showed that vSMCs provide baseline contractile tone, and that only depolarization of vSMCs results in alterations of vessel diameter and the regulation of CBF (Hill et al., 2015). It is important to note that hemodynamic variability within the microcirculation may account for some of these differences in findings (Gould et al., 2017). Overall, with the adult mouse model, there is no consensus on mural cell vasoactivity and the corresponding changes in CBF regulation.

Here we test the contractile properties of brain mural cells in zebrafish and determine the corresponding changes in cerebral vessel diameters during development. We find that vasoactivity changes dynamically during development. We show that both perictyes and vSMCs have the ability to actively regulate cerebral vessel diameters but this activity depends on the stage of development. Further, using live imaging and genetic ablation experiments, we show that pericytes on brain capillaries contract at earlier stages in development while brain vSMCs develop this ability later. However, once vSMCs have differentiated and matured, perictyes no longer regulate cerebral vessel diameter.

## METHODS

### Zebrafish husbandry and fish strains

All experimental procedures were approved by the University of Calgary’s Animal Care Committee (Protocol AC17-0189). Zebrafish embryos were maintained at 28.5°C and in E3 medium (Westerfield, 1995). Transgenic lines used include: *TgBAC*(*pdgfrβ:GFP*)^*ca41*^ (Whitesell et al., 2019), *TgBAC*(*pdgfrβ:Gal4*) ^*ca42*^(Whitesell et al., 2019), *Tg*(*kdrl:mCherry*) ^*ci5*^ (Proulx et al., 2010), *Tg*(*acta2:GFP*) ^*ca7*^(Whitesell et al., 2014), Tg(acta2:Gal4FF)^ca62^ (Whitesell et al., 2019), *Tg*(*flk:GFP*) ^*la116*^ (Choi et al., 2007), *Tg*(*UAS:NTR-mCherry*) ^*c264*^ (Davison et al., 2007), and *Tg*(*GFAP:GFP*)^*mi2001*^ (Bernardos and Raymond, 2006).

### Pharmacological agents and nitroreductase ablation

All chemicals were obtained from Sigma Aldrich (St. Louis, MO). Sodium nitroprusside dihydrate (SNP; 71778) was used at 1 mM. S-Nitroso-N-acetyl-DL-penicillamine (SNAP; N3398) was used at 100 μM. Diethylamine NONOate sodium salt hydrate (D184; NONOate) was used at 5 μM. (R)-(-)-Phenylephrine hydrochloride (P6126) was used at 10 μM. (-)-Norepinephrine (A7257) was used at 2 μM. The EP4 receptor antagonist, AH23848 hemicalcium salt hydrate (A8227), was used at 10 μM. All vasoactive agents were dissolved in E3 fish medium except for AH23848, which was dissolved in 0.1% DMSO in E3.

For ablation experiments, metronidazole (Mtz; M3761) was used at 50 μM at 3 dpf mural cell ablation. At 5 dpf, 50 μM was used for vSMC ablation and 1 mM for pericyte ablation.

### Heart rate measurements

A dose response curve was used to determine optimal drug concentrations. Heart rate was counted manually for a period of one minute. For each concentration of a vasoactive agent, 5-6 zebrafish were exposed individually to either E3 fish medium or the vasoactive agent.

### Microscopy and image analysis

Zebrafish were mounted on glass bottom petri dishes (MatTek, Ashland, MA, Cat. No. P50G-0-30-F), using 0.8% low melt agarose (Invitrogen (Carlsbad, CA) 16520-050), which was dissolved in E3 fish medium. Confocal imaging was conducted with a Zeiss LSM 700 confocal microscope. All images were obtained with the 488 nm and 555 nm lasers, with a pinhole of 1 airy unit (AU). All images were 12 bit with slice intervals of 1-2 μm, line step of 2, with a 10X (NA 0.25 Ph1) or 20X (NA 0.8) objective and either 512×512 or 1024×1024 in frame size. The z-stacks varied in range from 35 slices to 240 slices depending on the image. Images were processed using Zen Blue and or ImageJ/Fiji. In ImageJ/Fiji, maximum intensity projection was carried out on the confocal images for all images prior to any measurements.

Vessel diameters were measured at positions where there was an association of a mural cell with the endothelium. The vessel diameter was measured from the external diameter of the endothelium. Three measurements were obtained from each mural cell-endothelium association region and averaged to obtain a vessel diameter. All vessel diameter measurements were paired before and after treatment.

### Statistics

All data were graphed as normalized to baseline measurements unless otherwise indicated. GraphPad Prism7 was used to carry out all statistics using either a paired two-tailed t-test, unpaired t-test or repeated measures ANOVA with P≤ 0.05 = *, P≤ 0.005 = **, P≤ 0.0005 = ***. From the repeated measures ANOVAs, p values from Dunnett’s test are reported unless otherwise indicated. All data are represented as mean ± standard deviation (SD).

## RESULTS

### Pericytes and vSMCs appear on cerebral vessels of different diameters

Pericytes and vSMCs are first visualized in the zebrafish brain at 48 hpf and around 80 hpf respectively, and their abundance increases over time. We examine mural cell activity at two stages of development: 4 dpf, when vSMCs start to express *acta2*, and when *pdgfrβ* expressing pericytes have been present for about 2 days, and 6 dpf, when vSMCs have been present for about 2 days. We first identified stereotypical locations for pericytes and vSMCs. At 4 dpf, pericytes are present on smaller vessels with diameters ≤6.5 μm (Supplementary Figure 1 A and B), typically the brain Central Arteries (CtAs). vSMCs cover the larger vessels that are ≥9.5 μm in diameter (Supplementary Figure 1A and F). Early vSMC-covered vessels (at 4 dpf) include the caudal division of the internal carotid (CaDI) and the basilar artery (BA). Cerebral vessels with diameters between 6.5μm and 9.5μm have variable coverage by both pericytes and vSMCs and were excluded from further analysis as we could not clearly separate functions of mural cells in this size range of vessels.

In the zebrafish brain at 4 dpf pericytes appear sporadically along vessels with long processes that extend along the surface of the endothelium (Supplementary Figure 1C-E’’’). vSMCs appear on only a few brain vessels and form a continuous sheet around the vessel (Supplementary Figure 1G-I’’’). By 6 dpf, the total numbers of pericytes and vSMCs is higher than at 4 dpf. There are more pericytes than vSMCs at both stages (Figure 1A-B). At 6 dpf, vessels with diameters ≤6.5 μm continue to be primarily covered by pericytes while those with diameters of ≥9.5 μm are primarily covered by vSMCs. At 6 dpf, pericyte processes are in close proximity and appear to contact each other (Figure 1C-F’’’). vSMC coverage of the cerebral endothelium also extends to more dorsal vessels over time (Figure 1G-J’’’).

**Figure 1:**
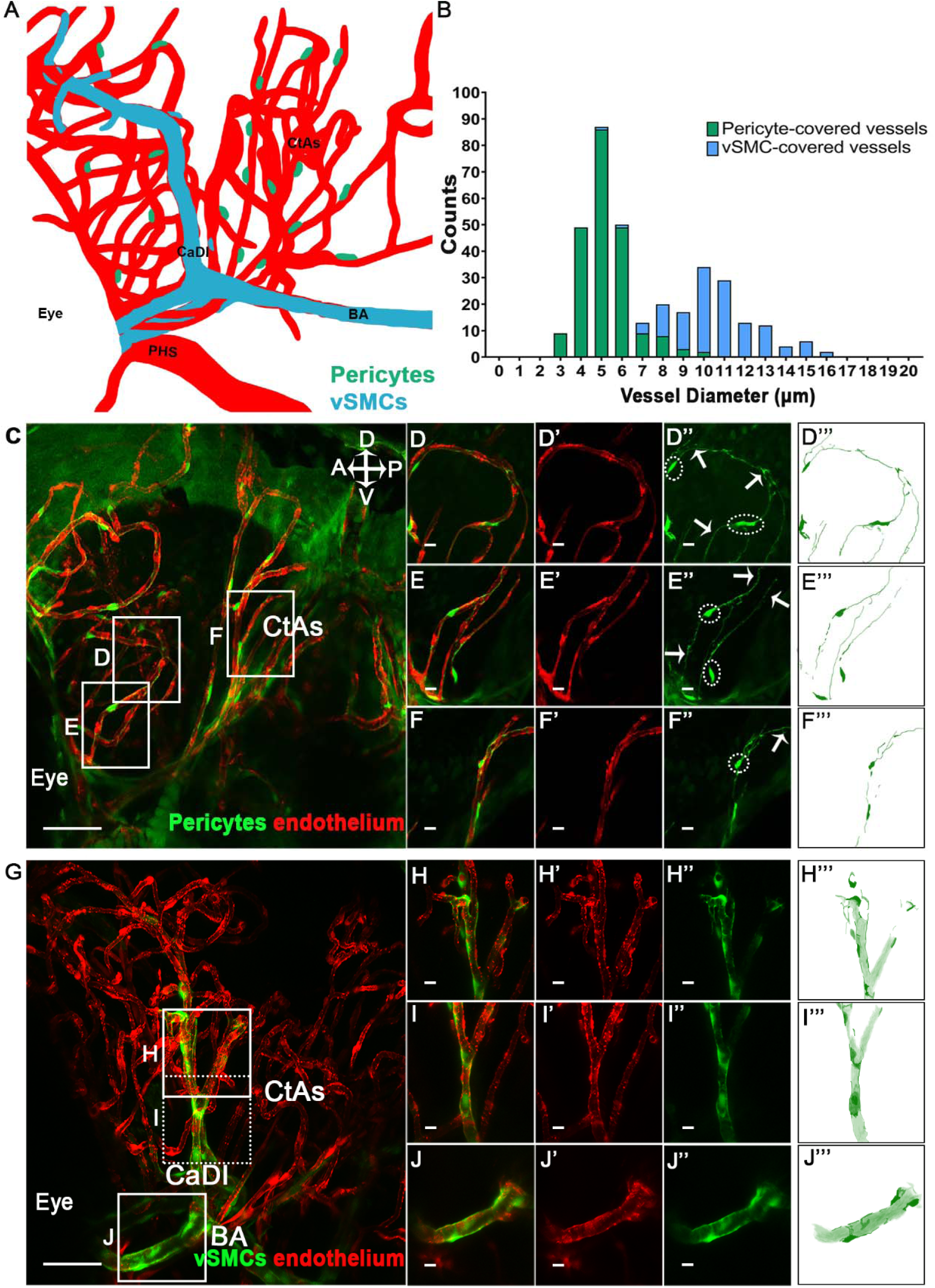
Mural cell morphology and coverage of the midbrain cerebral vasculature at 6 dpf. **A)** A model of vSMC and pericyte locations on the 6 dpf cerebral vessels highlighting the primordial hindbrain channel (PHS), internal carotid artery (CaDI), basilar artery (BA) and central arteries (CtAs). **B)** Graph of the number of cells counted on vessels of different diameters and expressing either a pericyte or vSMC transgene. **C)** Lateral image of cerebral vessel pericyte coverage at 6 dpf. **D-F’’’)** Enlargements of vessels in C highlighting contact between the processes of different pericytes. The circles highlight pericyte cell bodies while the arrows point to their processes. **G)** Lateral image of cerebral vessel vSMC coverage at 6 dpf, particularly on the caudal division of the internal carotid artery (CaDI) and the basilar artery (BA). **H-J’’’)** Enlargements of the vessels in G showing extensive vSMC coverage. Scale bars represent 50 µm for **C** and **G**, and 10 µm for **D-F’’’** and **H-J’’’**. A, P, V and D refer to anterior, posterior, ventral and dorsal.

### Early contraction of pericytes to noradrenaline is lost when vSMCs mature

To determine if mural cells are capable of contraction during early development, we utilized α_1_ adrenergic agonists. We established 2 μM as the most effective dose of noradrenaline (NA) at 6 dpf. As changes in heart rate might affect vascular dynamics, we determined that heart rate was unchanged with this dose (Supplementary Figure 2A). To determine changes in vessel diameter, the cerebral vasculature was imaged before dosing (0 min) and repeatedly every minute after NA treatment (1-4 minutes; Figure 2A). At 4 dpf, pericyte-covered vessels constrict in response to NA within the first minute of treatment (P≤ 0.0001, Figure 2B, and D-H). Constriction persists for the duration of NA treatment and over time, vessels constrict further. The average pericyte-covered vessel diameters at 0, 1, 2, 3, and 4 minutes were 5.3 μm ± 0.7, 5.1 μm ± 0.7, 5.0 μm ± 0.7, 5.0 μm ± 0.8 and 5.0 μm ± 0.8 respectively. We note that of the 60 vessels studied, 31 vessels constrict consistently throughout the experimental period whereas the other pericyte-covered vessels behave inconsistently and demonstrate both dilation and constriction. Surprisingly, we find that at 6 dpf, pericyte-covered vessels no longer constrict to NA (Figure 2N). The average pericyte-covered vessel diameter at 0, 1, 2, 3 and 4 minutes were 5.1 μm ± 0.8, 5.2 μm ± 0.9, 5.0 μm ± 0.9, 5.2 μm ± 0.8 and 5.1 μm ± 0.8 respectively.

**Figure 2:**
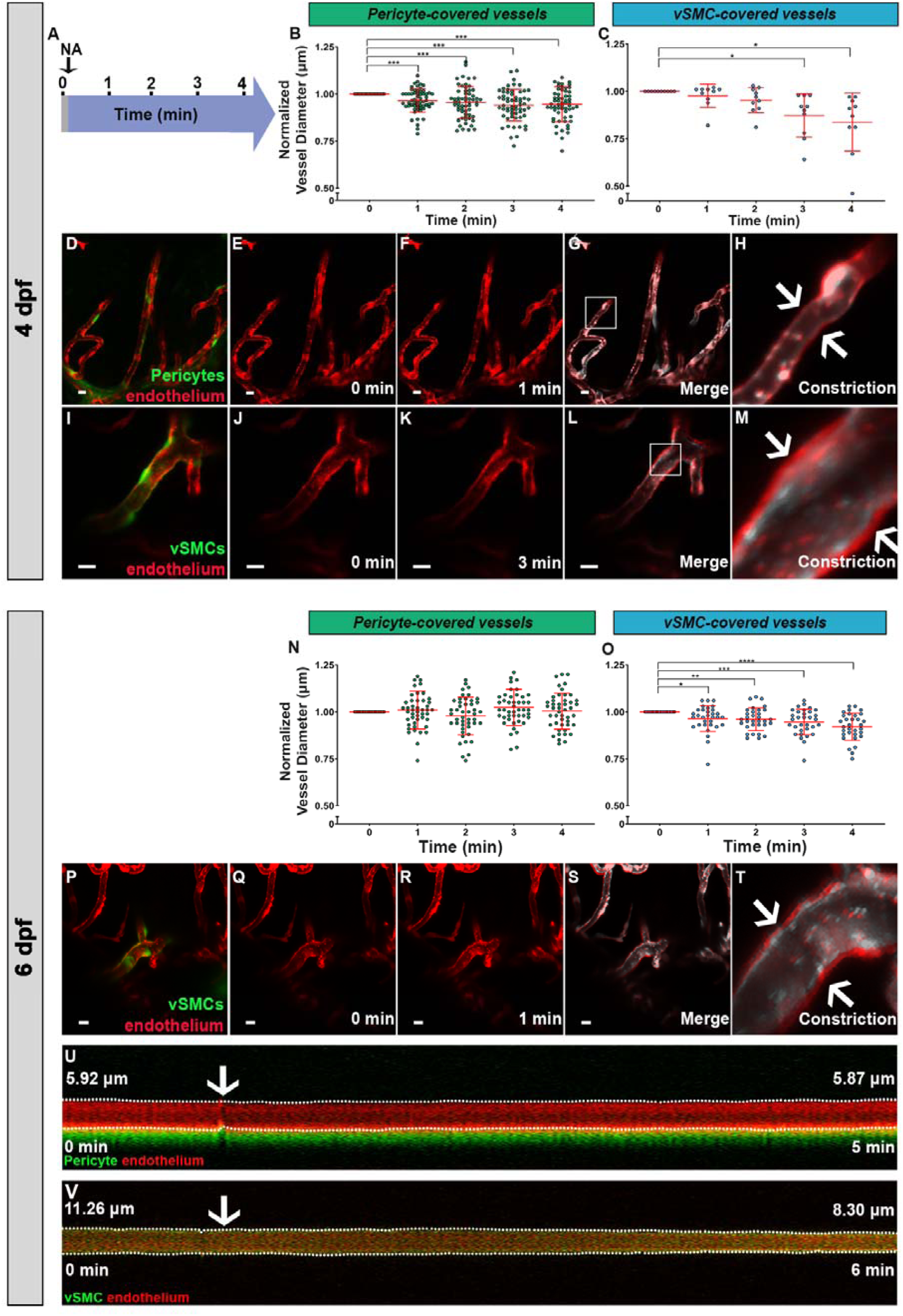
Vessel constriction in response to NA by both pericytes and vSMCs at 4 dpf, but only vSMCs at 6 dpf. **A)** Schematic of the experimental timeline. Imaging occurs prior to addition of 2 µM NA (0 min) and at one-minute intervals over 4 minutes after drug addition. **B)** At 4 dpf, pericyte-covered vessels constrict in response to NA within the first minute (P≤ 0.0001; n=60 measurements from 14 embryos). **C)** At 4 dpf, vSMC-covered vessels constrict within 3 minutes of treatment (P≤ 0.006; n=10 measurements from 9 embryos). **D-H)** Images of pericyte-covered vessel constriction. **I-M)** Images of vSMC-covered vessel constriction. **N)** At 6 dpf, pericyte-covered vessels do not constrict in response to NA (P≤ 0.019; n=46 measurements from 15 embryos). **O)** At 6 dpf, vSMC-covered vessels constrict in response to NA (P≤ 0.0001; n=32 measurements from 25 embryos). **P-T)** Images of vSMC-covered vessel constriction at 6 dpf. **U)** Kymograph showing the lack of diameter change over time from a pericyte covered vessel over a 5-minute period. **V)** Kymograph of a vSMC-covered vessel constricting over a 6-minute period. Arrow marks the start of NA treatment in U and V. Scale bars on **D-G, I-L**, and **P-S** represent 10 µm. Significance was determined by ANOVA with P≤ 0.05 = *, P≤ 0.005 = ** and P≤ 0.0005 = ***.

Larger vSMC-covered vessels also constrict in response to NA at 4 dpf, typically after 3 minutes of exposure (P≤ 0.006, Figure 2C, and I-M). The average vSMC-covered vessel diameters at 0, 1, 2, 3, and 4 minutes were 11.5 μm ± 1.2, 11.3 μm ± 1.5, 11.0 μm ± 1.4, 10.0 μm ± 1.6, and 9.6 μm ± 1.8 respectively. All measured vessels constrict by 3 minutes and remain constricted. At 6 dpf, the vessels constrict faster than they did at 4 dpf, responding within the first minute, and remaining significantly constricted for the duration of drug exposure (P≤ 0.0001, Figure 2O and P-T). The average vSMC-covered vessel diameters at 0, 1, 2, 3 and 4 minutes were as follows: 11.8 μm ± 1.9, 11.3 μm ± 2.0, 11.2 μm ± 1.7, 11.1 μm ± 1.7, and 10.8 μm ± 1.8. Kymographs show that over time, a pericyte-covered vessel does not significantly decrease in diameter (Figure 2U), while a vSMC-covered vessel decreases in diameter (Figure 2V).

Thus, both pericyte and vSMC-covered vessels constrict at 4 dpf, with pericytes responding faster. However, at 6 dpf, only vSMC contraction occurs and it occurs faster relative to 4 dpf, suggesting that vSMCs have differentiated further. However, the response of vSMCs is variable; of 32 measured vessels at 6 dpf, only 13 constrict consistently over the experimental period, suggesting that myogenic response is still maturing in many vSMCs.

### Early contraction of pericytes to phenylephrine is lost when vSMCs mature

We used a second α_1_ adrenergic agonist, phenylephrine (PE) to test whether it would cause a similar contractile behavior in mural cells. A dose response curve identified 10 µM as the optimal dose at 6 dpf, with no effect on heart rate (Supplementary Figures 2B). Similar to NA administration, pericyte-covered vessels constrict in response to PE at 4 dpf but not at 6 dpf (Figure 3). At 4 dpf, the average pericyte-covered vessel diameters at 0, 1, 2, 3, and 4 minutes were 5.2 µm ± 0.9, 5.2 µm ± 0.8, 5.1 µm ± 0.9, 5.1 µm ± 0.9, 5.1 µm ± 0.8 and 5.0 ± µm 0.8 respectively. These vessels constrict within 3 minutes and maintained this to 5 minutes (P≤ 0.007, Figure 3B and C-G). Also, like NA, only a sub-population of pericyte-covered vessels constrict consistently (35 of 55 vessels constrict at both 3 and 5 minutes). Similar to NA at 6 dpf, pericyte-covered vessels do not constrict to PE at 6 dpf (Figure 3H). At 6 dpf, the average pericyte-covered vessel diameters at 0, 1, 2, 3, 4 and 5 minutes were 4.7 µm ± 0.8, 4.7 µm ± 0.8, 4.7 µm ± 0.7, 4.7 µm ± 0.8, 4.8 µm ± 0.8 and 4.7 µm ± 0.9 respectively.

**Figure 3:**
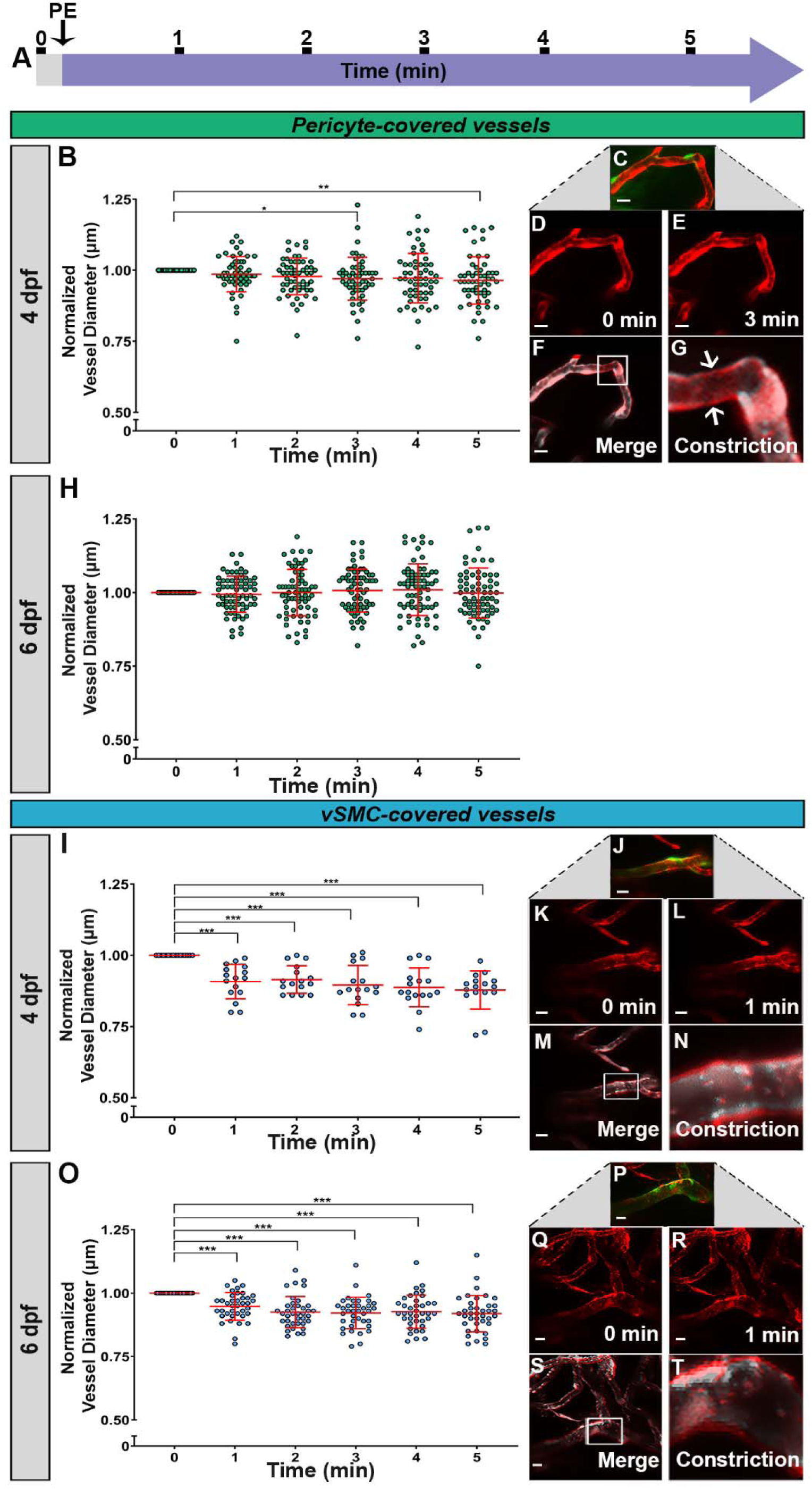
Vessel constriction in response to PE by both pericytes and vSMCs at 4 dpf, but only vSMCs at 6 dpf. **A)** Schematic of the experimental timeline. Imaging occurs prior to addition of 10 µM PE (0 min) and at one-minute intervals over 5 minutes after drug addition. **B)** At 4 dpf, pericyte-covered vessels constrict in response to PE (P≤ 0.007; n= 55 measurements from 15 embryos). **C-G)** An example of pericyte-covered vessel constriction at 4 dpf. **H)** Pericyte-covered vessels do not constrict at 6 dpf (P≤ 0.552; n=68 measurements from 15 embryos). **I)** vSMC-covered vessels constrict within the first minute of PE treatment at 4 dpf (P≤ 0.0001; n=16 paired measurements from 15 embryos). **J-N)** An example of vSMC-covered vessel constriction at 4 dpf. **O)** At 6 dpf, vSMC-covered vessels constrict in response to PE (P≤ 0.0001; n= 38 paired measurements from 22 embryos). **P-T)** An example of the vSMC-covered vessel constriction at 6 dpf. Scale bars represent 10 µm in **C-F, J-M**, and **P-S**. Significance was determined by ANOVA with P≤ 0.05 = *, P≤ 0.005 = ** and P≤ 0.0005 = ***.

vSMCs constrict consistently at both 4 and 6 dpf in response to PE (Figure 3). At 4 dpf, vSMC-covered vessels constrict within the first minute to PE (P≤ 0.0001, Figure 3I-N), which is faster than the pericyte-covered vessels. The response of vSMCs to PE is also faster than their response to NA at 4 dpf (Figure 2C). At 4dpf, vSMC-covered vessel diameters at 0, 1, 3, 4 and 5 minutes were 12.0 µm ± 1.4, 10.9 µm ± 1.1, 11.0 µm ± 1.0, 10.7 µm ± 0.9, 10.6 µm ± 1.0 and 10.5 µm ± 1.0 respectively. These vessels constrict to a similar extent for the duration of PE exposure (14/16 vessels). At 6 dpf, vSMC-covered vessels constrict and their average vessel diameters at 0, 1, 2, 3, 4 and 5 minutes were 11.7 µm ± 1.6, 11.0 µm ± 1.3, 10.8 µm ± 1.5, 10.7 µm ± 1.6, 10.8 µm ± 1.5 and 10.7 µm ± 1.6 respectively (P≤ 0.0001, 27/38 vessels, Figure 3O-T).

Thus, each mural cell responds similarly to each of the vasoconstrictors in this study (NA and PE). At 4 dpf, both pericytes and vSMCs contract in response to these agents whereas at 6 dpf, only the vSMCs contract. This highlights the activity of pericytes at the earlier stage, and the dominance of vSMCs at the later stage.

### Pericytes and vSMCs actively constrict cerebral vessels

Genetic ablation of specific lineages was used to determine which mural cells actively regulate cerebral vessel diameter. We utilized transgenic lines expressing Gal4 under the *pdgfrβ* (Ando et al., 2016) or acta2 promoters (Whitesell et al., 2014) for pericytes and vSMCs respectively. Selective ablation of the mural cell lineage was possible by crossing transgenic lines expressing the nitroreductase gene under a UAS promoter and exposing embryos to Mtz for 24 hours. After ablation, zebrafish were exposed to vasoconstricting agents to determine changes in vessel diameter. Ablation started at 3 dpf for 4 dpf measurements, and at 5 dpf for 6 dpf experiments. The fish had normal morphology (Supplementary Figure 3A – H) when mural cells were ablated (Supplementary Figure 3A’-H’).

At 4 dpf we found constriction of cerebral vessels was reduced after ablation. At 4 dpf, after pericyte ablation, cerebral vessels with diameters of ≤6.5 μm (normally vessels with pericyte coverage), still constrict in response to NA (P≤ 0.001, Supplementary Figure 4B, and F-I), although they constrict later than in the unablated fish (Figure 2B). The vessel diameters at 0, 1, 2, 3 and 4 minutes were 5.6 µm ± 0.5, 5.6 µm ± 0.6, 5.5 µm ± 0.6, 5.4 µm ± 0.5 and 5.3 µm ± 0.5 respectively. These ≤6.5 μm diameter vessels with ablated pericytes also constrict in response to PE (P≤ 0.0003, Supplementary Figure 4C and J-M). The average vessel diameters at 0, 1, 2, 3, 4 and 5 minutes were 5.3 µm ± 0.8, 5.2 µm ± 0.8, 5.1 µm ± 0.8, 5.1 µm ± 0.8, 5.1 µm ± 0.8, and 5.01 µm ± 0.8 respectively. The vessel diameters significantly decreased at 3 minutes post treatment and remained constricted. We hypothesized that incomplete ablation might account for remaining contractile activity, and therefore assessed ablation efficiency of cerebral pericytes or vSMCs at 4 dpf. We note partial ablation of both lineages (Supplementary Figure 3I-N’’). As the dose of Mtz used for the ablation was the maximal tolerated dose at this developmental stage without inducing morphological defects, the observation of reduced contractile ability is consistent with pericytes playing an active role in vascular constriction at 4 dpf.

Similarly, cerebral vessels normally covered by vSMCs constrict in embryos with incompletely ablated vSMCs at 4 dpf but it was weaker and not maintained. Cerebral vessels with diameters of ≥9.5 µm, normally covered with vSMCs constrict in response to NA with ablated vSMCs (P≤ 0.001, Supplementary Figure 4D and N-Q) and remain constricted for the duration of NA treatment. The average vessel diameter of these vSMC ablated vessels at 0, 1, 2, 3 and 4 minutes were 11.6 µm ± 1.4, 11.0 µm ± 1.8, 10.3 µm ± 1.7, 9.7 µm ± 1.9, and 9.5 µm ± 2.1 respectively. These ≥9.5 µm in diameter vessels also constrict in response to PE when vSMCs are ablated (P≤ 0.070, Supplementary Figure 4E, and R-U). The average vSMC-covered vessel diameters at 0, 1, 2, 3, 4 and 5 minutes were 11.7 µm ± 2.1, 11.3 µm ± 2.1, 11.3 µm ± 2.0, 11.3 µm ± 2.1, 11.2 µm ± 2.01 and 11.2 µm ± 2.0 respectively. Significant constriction only occurred between 0 and 4 minutes but interestingly, these vessels did not remain constricted. Thus, constriction in embryos with ablated vSMCs is later and not maintained (Figure 3I). This reduction in contractile ability is consistent with active contraction by vSMCs at 4 dpf.

When mural cells are ablated at 5 dpf and tested at 6 dpf, we see a complete reduction of contractile activity as neither pericyte nor vSMC covered vessels constrict (Figure 4). Cerebral vessels with ≤6.5 in diameter no longer respond to either NA or PE (Figure 4B-C). The average vessel diameter at 0, 1, 2, 3 and 4 minutes with NA treatment were 4.5 µm ± 0.9, 4.4 µm ± 0.9, 4.4 µm ± 0.9, 4.5 µm ± 1.0 and 4.5 µm ± 1.0 respectively. As for PE treatment, the average vessel diameters at 0, 1, 2, 3, 4 and 5 minutes were 4.8 µm ± 0.8, 4.9 µm ± 0.7, 4.9 µm ± 0.8, 4.9 µm ± 0.8, 4.9 µm ± 0.8 and 4.8 µm ± 0.8 respectively. Similarly, ≥9.5 µm diameter vessels with ablated vSMCs do not constrict to either NA or PE at 6 dpf (Figure 4D-E). The average vessel diameters at 0, 1, 2, 3 and 4 minutes of treatment with NA were 13.5 µm ± 2.9, 13.5 µm ± 2.7, 13.3 µm ± 3.0, 13.2 µm ± 3.0 and 13.1 µm ± 3.1 respectively. After treatment with PE, the average vessel diameters at 0, 1, 2, 3, 4 and 5 minutes were 12.5 µm ± 2.1, 12.4 µm ± 2.5, 12.4 µm ± 2.7, 12.1 µm ± 2.3, 12.2 µm ± 2.3 and 12.1 µm ± 2.4 respectively. Our 6 dpf genetic ablation data suggest that vSMCs actively contract at 6 dpf to regulate vessel diameter.

**Figure 4:**
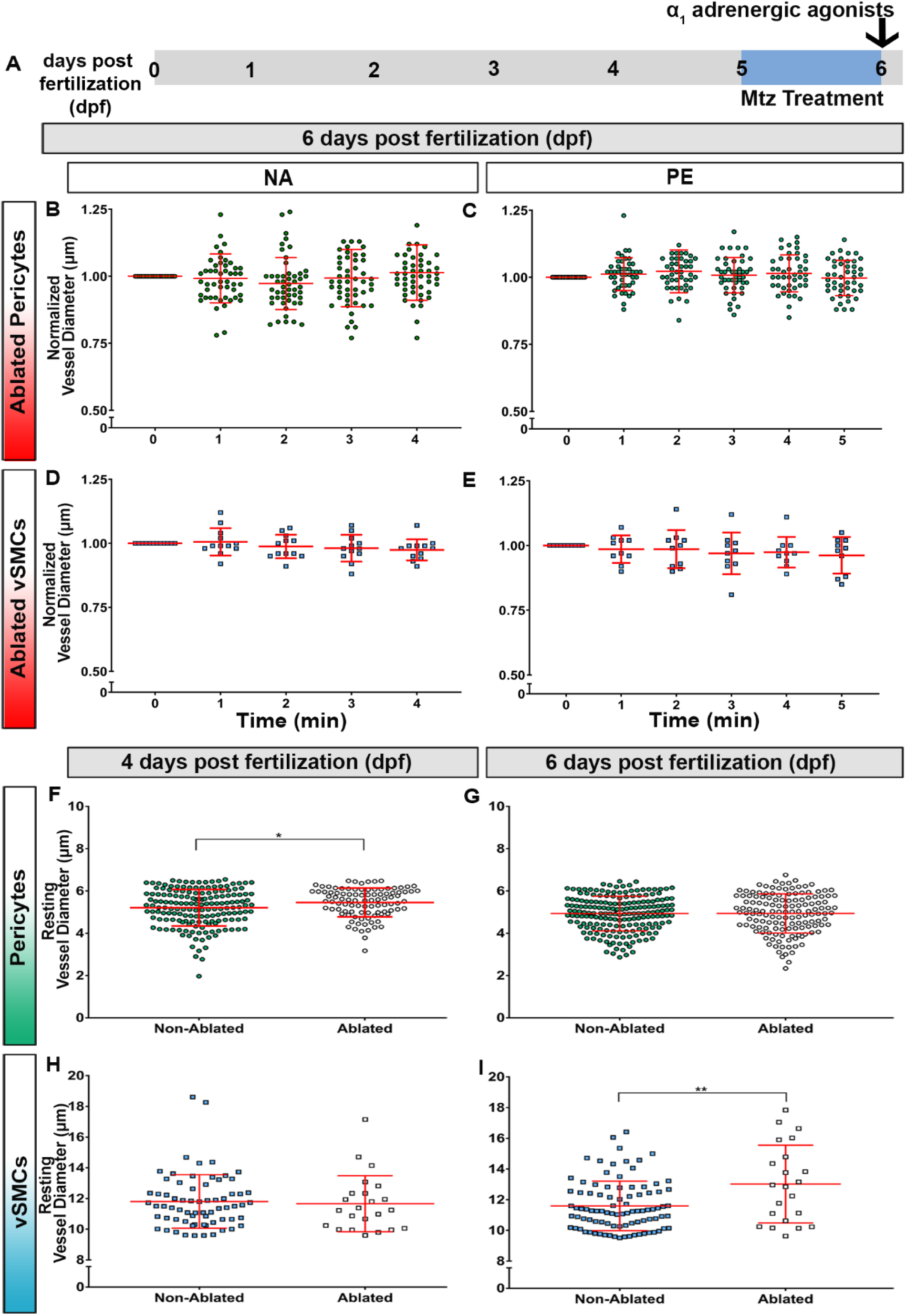
Lack of cerebral vessel constriction in the absence of mural cells at 6 dpf. **A)** Experimental timeline. Embryos were treated with Mtz from 5-6 dpf before treatment with vasoconstrictors. Vessels with ablated pericytes (diameters of ≤6.5 μm) did not constrict to either drug, **B)** NA (P≤ 0.065, n=50 measurements from 12 embryos) or **C)** PE (P≤ 0.123, n=45 measurements from 6 embryos). Vessels with ablated vSMCs (diameters of ≥9.5 µm) also did not constrict to either **D)** NA (P≤ 0.233, n=12 measurements from 10 embryos) or **E)** PE (P≤ 0.314, n=10 measurements from 9 embryos) after vSMC ablation. **F)** Furthermore, vessels with ablated pericytes have an enlarged diameter at 4 dpf (P≤ 0.016, n=168 and 97 for the non-ablated and ablated groups). **G)** However, there is no difference in vessel diameter in the presence and absence of pericytes at 6 dpf (P ≤0.953, n=193 and 141 for the non-ablated and ablated groups). **H)** There is no difference in vessel diameter in the presence and absence of vSMCs at 4 dpf (P≤ 0.738, n=68 and 23 for the non-ablated and ablated groups). **I)** However, vessels with ablated vSMCs have an enlarged diameter at 6 dpf (P≤ 0.001, n=92 and 22 for the non-ablated and ablated groups). Significance was determined by repeated measures one-way ANOVA for **B-E** and unpaired two-tailed t-tests for **F-I** with P≤ 0.05 = * and P≤ 0.005 = **.

Additionally, we used another method to determine whether pericytes and vSMCs actively regulate vascular tone by comparing resting vessel diameter in the presence and absence of each mural cell. Vascular tone is the active regulation of a constant resting vessel diameter that is less than maximal dilation. At 4 dpf, vessels with ablated pericytes are significantly enlarged (P≤ 0.016, Figure 4F). The average vessel diameter with pericytes present was 5.2 µm ± 0.9, while without pericytes it was 5.5 µm ± 0.7. At 6 dpf as expected, the average vessel diameter in the presence and absence of pericytes remain unchanged, 4.9 µm ± 0.8 and 4.9 µm ± 0.9 (Figure 4G), which is in agreement with our pharmacological data suggesting that pericytes regulate vessel diameter at 4 dpf but not 6 dpf.

vSMCs regulate tone in a complementary manner. At 4 dpf, there is no difference in cerebral vessel diameter in the presence and absence of vSMCs (Figure 4H). The average vessel diameter with and without vSMCs was 11.8 µm ± 1.8 and 11.7 µm ± 1.8 respectively. However, at 6 dpf, the average vessel diameter with vSMC coverage was 11.6 µm ± 1.6, while with ablated vSMCs it was 13.0 µm ± 2.5 (P≤ 0.001, Figure 4I).

Taken together, we provide evidence that at 4 dpf, pericytes are a key regulator of cerebral vessel diameter, but at 6 dpf, vSMCs become the primary regulators of vascular tone.

### Vessel dilation is mediated by pericytes and vSMCs

Nitric oxide is a key molecule mediating vascular mural cell relaxation. To test the ability of developing mural cells to relax, we exposed zebrafish to nitric oxide donors at different stages of development. Dose response curves were carried out for each agent and changes to heart rate were assessed and showed no effect (Supplementary Figure 2C). At 4 dpf, pericyte-covered vessels dilate in response to SNP (P≤ 0.005; Figure 5A and E-I). The average pericyte-covered vessel diameter before and after SNP treatment was 5.1 µm ± 1.0 and 5.4 µm ± 1.1 respectively (P≤ 0.010, 33/53 vessels, Figure 5B). To determine if this dilation is due to the active relaxation of pericytes, we genetically ablated pericytes with Mtz and then exposed the zebrafish to SNP. After ablation, vessels with diameters ≤6.5 µm constricted instead of dilating (P≤ 0.003, Figure 5C), from 5.7 µm ± 0.5 to 5.4 µm ± 0.5 in response to SNP (P≤ 0.001, 25/33 vessels, Figure 5D). The loss of dilation in the absence of pericytes supports the active role of pericytes in vascular dilation at 4 dpf.

**Figure 5:**
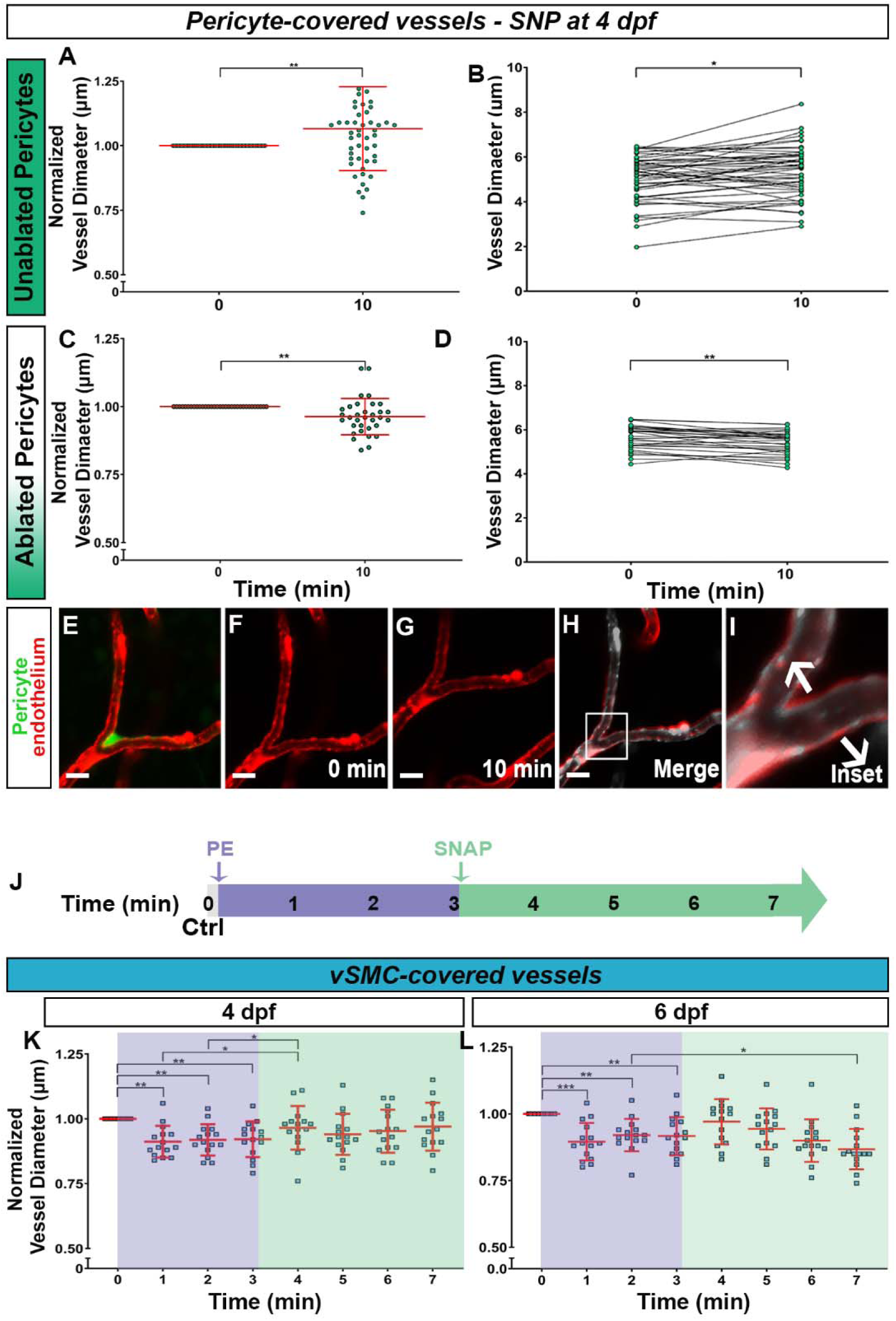
Pericyte-covered vessels dilate at 4 dpf while vSMC dilate at 6 dpf, but only from a pre-constricted state. **A)** Pericyte-covered vessels dilate in response to SNP at 4 dpf (P≤ 0.005, n=53 measurements from 15 embryos). **B)** 33/53 pericyte-covered vessels dilate (P≤ 0.010, n =53 measurements from 15 embryos). **C)** When pericytes are ablated, vessels with diameters of ≤6.5 μm constrict in response to SNP at 4 dpf (P≤ 0.003, n=33 measurements from 10 embryos). On average the vessel diameters decreased from 5.65 µm to 5.43 µm (P≤ 0.001, **D). E-I)** An example of pericyte-covered vessel dilation to SNP at 4 dpf. **J)** Experimental timeline for the pre-constriction followed by dilation of vSMC-covered vessels. Embryos were imaged prior to drug addition (0 min), followed by 3 minutes of exposure to PE (10 µM) and then to 4 minutes of exposure to SNAP (100 µM). Embryos were imaged every minute. **K)** At 4 dpf, vSMC-covered vessels do not dilate from a pre-constricted state (P≤0.0001, n=15 measurements from 11 embryos) after SNAP exposure. **L)** At 6 dpf, vSMC-covered vessels dilate from a pre-constricted state (P≤ 0.0002, n=15 measurements from 13 embryos). The shading in K and L highlight the drug exposures and the corresponding changes in vSMC-covered vessel diameter according to the experimental timeline **(J).** Scale bars represent 10 µm for **E-I** and significance was determined by paired two-tailed t-tests with P≤ 0.05 = *, P≤ 0.005 = ** and P≤ 0.0005 = ***.

We also assessed the relaxation of vSMCs to SNP at 4 dpf, however surprisingly, there was no significant dilation (Supplementary Figure 5A). The average vSMC-covered vessel diameter before and after SNP treatment was 11.8 µm ± 2.0 and 11.8 µm ± 3.0 (Supplementary Figure 5B). We noted a binary response from vSMC-covered vessels. Of 42 measurements, 16 dilated, one did not change in diameter while 25 constricted (Supplementary Figure 5B). This suggests that at 4 dpf, the ability of vSMCs to relax may still be maturing.

We reasoned that if a vessel is close to maximal dilation, it may not be able to show a large dilation. Thus, we tested whether vessels are able to dilate from a pre-constricted state. We contracted vSMCs using 10 µM of PE and then exposed them to 100 µM nitric oxide donor SNAP (Figure 5J). Similar to our previous results at 4 dpf, vSMCs contracted but were unable to relax (P≤ 0.0001, Figure 5K). The vSMC-covered vessels constricted to PE within the first minute (and remained constricted for the duration of PE exposure, 2 minute and 3-minute, P ≤0.003 and P ≤0.008 accordingly). The average vessel diameters at 0, 1, 2 and 3 minutes were 12.3 µm ± 1.8, 11.0 µm ± 1.7, 11.3 µm ± 1.1, and 11.2 µm ± 1.2 respectively. Some vessels appear to dilate in response to SNAP within first minute of treatment (4^th^ minute), however, as time progresses, they constrict again and remain constricted for the duration of the experiment. The vessel diameters at 4, 5, 6 and 7 minutes were 11.9 µm ± 1.2, 11.5 µm ± 1.0, 11.0 µm ± 1.0 and 10.6 µm ± 1.4 respectively. Thus, we conclude that at 4 dpf, vSMC-covered vessels only constrict but do not dilate.

In contrast, when vSMCs are more mature at 6 dpf, they can relax from a pre-contracted state (P≤ 0.0002, Figure 5L). In response to PE, vSMC covered vessels constrict (P≤ 0.001 between 0 and 1 minutes, P≤ 0.002 between 0 and 2 minutes and P≤ 0.010 between 0 and 3 minutes). The average vessel diameter at 0, 1, 2, and 3 minutes were 11.1 µm ± 0.8, 10.1 µm ± 1.0, 10.2 µm ± 0.9 and 10.2 µm ± 1.0 respectively. When SNAP was added, these vessels then actively dilate (4^th^ minute). There was significant dilation from experimental minute 1 (10.1 µm ± 1.0) to minute 4 (10.7 µm ± 1.1) (P≤ 0.035). Thus by 6 dpf, vSMCs have developed active myogenic ability to both contract and relax.

### Pericytes relax via the EP4 receptor to cause vessel dilation

Astrocyte endfeet release prostaglandins (including PGE2, the ligand of EP4 receptor) to evoke myogenic responses in pericytes (Macvicar and Newman, 2015). In the mouse, pericytes relax via prostaglandin EP4 receptors (Hall et al., 2014). Blocking the EP4 receptor using the antagonist AH23848 is predicted to constrict vessels. When this is followed by inducing dilation using SNAP (Figure 6A) we can test the role of EP4 receptors in pericyte relaxation. In response to AH23848 at 4 dpf, pericyte-covered vessels constrict within the first minute and remain constricted (P≤ 0.002, Figure 6B and D-H). The average vessel diameters at 0, 1, 2, 3 and 4 minutes of exposure to AH23848 were 5.4 µm ± 0.7, 5.3 µm ± 0.7, 5.3 µm ± 0.7, 5.2 µm ± 0.8, and 5.1 µm ± 0.8 respectively. After SNAP addition, these vessels remained constricted. The average vessel diameter at 5, 6, 7 and 8 minutes were 5.1 µm ± 0.9, 5.2 µm ± 0.8, 5.2 µm ± 0.8 and 5.3 µm ± 0.8 respectively. Thus at 4 dpf, blocking the EP4 receptor/pathway constricts pericyte-covered vessels and compromises the ability of pericytes to respond to NO donor SNAP.

**Figure 6:**
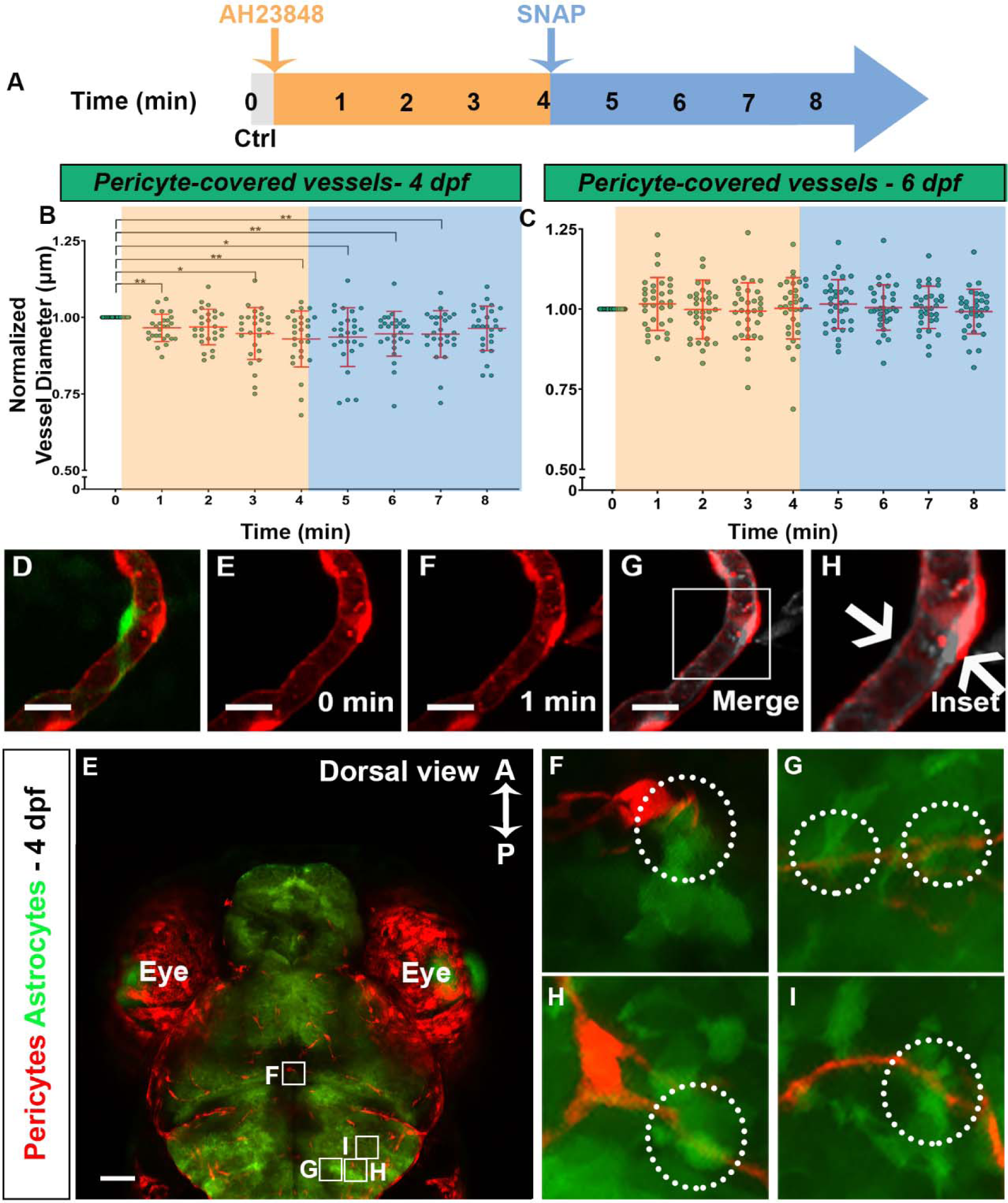
Pericytes relax via the EP4 receptor to cause vessel dilation. **A)** Experimental timeline. The embryos were imaged prior to drug treatment to obtain baseline vessel diameter (0 min). AH23848 was then added and embryos were imaged every minute for four minutes. SNAP was added after 4 minutes, and the fish were imaged every minute for the next four minutes. **B)** At 4 dpf, pericyte-covered vessels constrict in response to AH23848 and remain constricted after SNAP treatment (P≤ 0.002, n= 27 measurements from 9 embryos). **C)** At 6 dpf, the pericyte-covered vessels do not respond to either vasoactive agent (P≤ 0.55, n= 33 measurements from 15 embryos). The shading in B and C reflect drug exposure windows as in the experimental timeline. **D-H)** An example of pericyte-covered vessel constriction in response to AH23848 at 4 dpf. **E)** Spatial relationship between pericytes and astrocytes (marked with GFAP:GFP) at 4 dpf. A and P refer to anterior and posterior. **F-I)** Enlargements of D. White circles in F-I highlight close contact between the two cell types. Scale bars represent 50 µm in **E** and 10 µm in **F-I**. Significance was determined by one-way ANOVA with P≤ 0.05 = *.

Similar to our previous data at 6 dpf, we found AH23848 has no effect on pericyte-covered vessels (P≤ 0.55, Figure 6C). As expected, these vessels also do not dilate in response to SNAP after treatment with AH23848. At 6 dpf, the average vessel diameters at 0, 1, 2, 3, 4, 5, 6, 7 and 8 minutes were 4.4 µm ± 0.6, 4.5 µm ± 0.6, 4.4 µm ± 0.6, 4.4 µm ± 0.6, 4.4 µm ± 0.6, 4.5 µm ± 0.7, 4.4 µm ± 0.6, 4.4 µm ± 0.6 and 4.4 µm ± 0.6 respectively.

It was important to determine the spatial relationship between pericytes and astrocytes at 4 dpf since both cell types are part of the neurovascular unit. We observed direct contact between pericytes and astrocytes in the 4 dpf zebrafish brain (Figure 6E-I), suggesting that astrocytic regulation of pericyte behavior via prostaglandins would be possible at this stage (Figure 6E-I). Taken together, our pharmacological data showing regulation of pericytes via prostaglandin receptor and the physical proximity between pericytes and astrocytes, this is consistent with pericytes responding to prostaglandin signals at this stage to regulate vessel diameter.

## DISCUSSION

### Vasoactive regulation in early zebrafish development

The advantages of optical clarity and availability of transgenic lines for pericyte and vSMC lineages in zebrafish allow us to explore the dynamic role of mural cells in maintaining vascular tone during embryonic development. We have done so for the first time using live imaging. No studies have been conducted previously during embryonic development, as development of vascular myogenesis occurs in utero in mammals and cannot be live imaged. The roles of pericytes and vSMCs in regulating vessel diameter are conflicting in the literature for several reasons; unlike in zebrafish, markers such as *pdgfr*β can label both mural cell types in mouse (Armulik et al., 2011; Nakayama et al., 2013), and mural cells have also been identified based on morphology, which can be unreliable (Attwell et al., 2016). Pericytes also have a variety of morphologies (Hartmann et al., 2015); some pericytes express contractile proteins and some do not. These factors make it difficult to clearly separate mural cell contribution to vascular tone regulation, potentially leading to contradictory conclusions regarding myogenic activity (Fernandez-Klett et al., 2010; Hill et al., 2015).

We show a switch in mural cell types that regulate cerebral vessel diameters at different stages of development. As the cerebral vasculature develops, mural cell numbers increase from 4 dpf to 6 dpf. At 4 dpf, pericytes actively constrict and dilate cerebral vessels to maintain vascular tone. At this stage, vSMCs have the capacity to contract in response to vasoconstricting agents. However, the response is slow and there is heterogeneity in responses to the stimulus. At 6 dpf, vSMCs are the only cells we detect in the zebrafish cerebral vasculature that regulate vascular tone in our assays using pharmacological stimuli (Figure 7).

**Figure 7:**
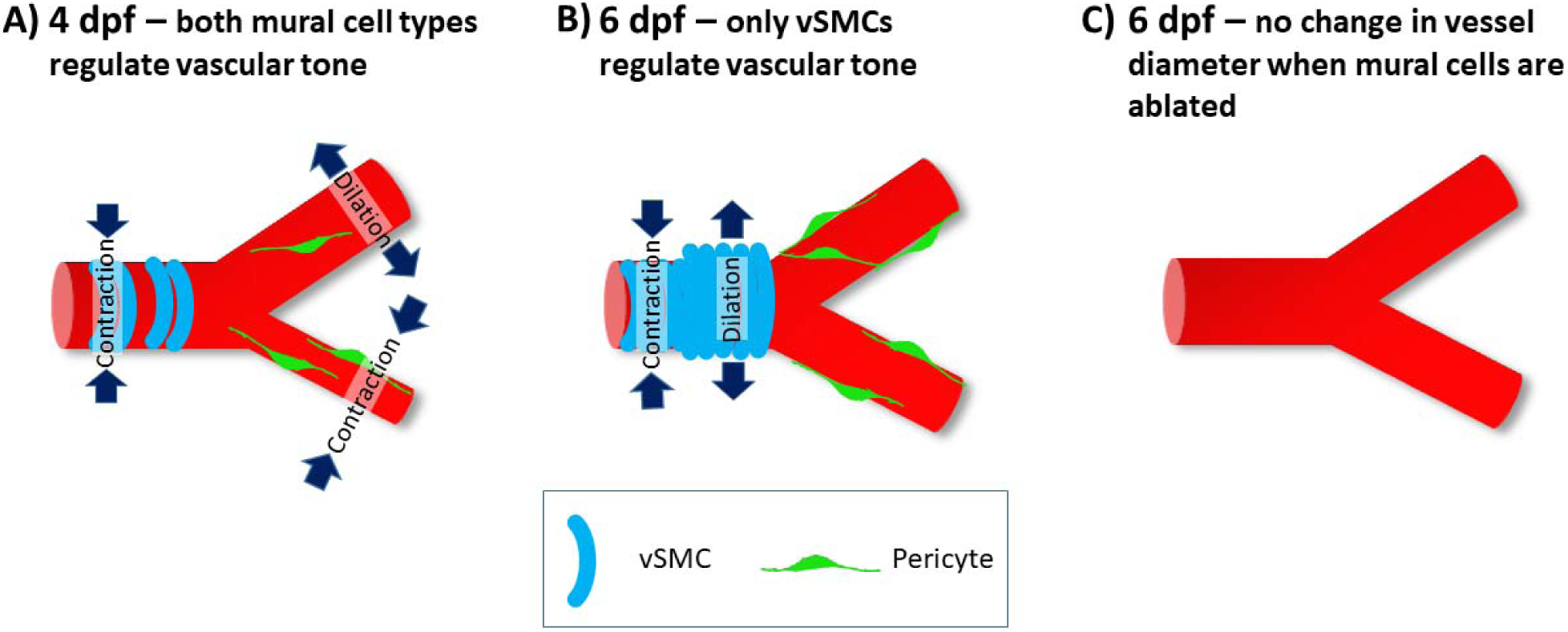
Summary Figure – Developmental changes in mural cell ability to regulate cerebral vascular tone. **A)** At 4 dpf, both pericyte and vSMC-covered vessels constrict while only pericyte-covered vessels dilate. **B)** At 6 dpf, both pericyte and vSMC populations have increased in number. Pericyte-covered vessels no longer respond to vasoactive agents while vSMC-covered vessels constrict but can only dilate from a pre-constricted state. **C)** There is no myogenic response when mural cells are ablated at 5 dpf and exposed to vasoactive agents at 6 dpf.

We find it intriguing that at 6 dpf, while vSMCs can contract, they can only relax from a precontracted state, suggesting that larger brain vessels in the zebrafish embryo are maintained in a dilated state during normal development. Interestingly, as hemodynamic load increases in the zebrafish trunk between 24-48 hpf, the diameter of the DA and posterior cardinal vein (PCV) increases (Sugden et al., 2017). However, by 72 hpf, endothelial cells align themselves by changing shape to decrease vessel diameter (Sugden et al., 2017). These time points align with pericyte and vSMC development and recruitment to the vessels. Once recruited to the vessels, these mural cells then regulate changes in vessel diameter by actively contracting or relaxing.

### Pericytes actively contract and relax to regulate cerebral vessel diameters

We provide several lines of evidence to show pericytes actively regulate vessel diameter. Firstly, our results show that at 4 dpf, pericytes are responsive to vasoactive agents, including vasoconstrictors (NA, PE and AH23848) and the vasodilator SNP. Secondly, genetic ablation of pericytes reduces vasoconstriction and removes vasodilation. Thirdly, the resting diameter of vessels with ablated pericytes is larger than wild type pericyte-covered vessels suggesting that pericytes contribute to the regulation of vessel diameter at this stage.

We observed a high, but not complete level of ablation of both pericytes and vSMCs at 4 dpf, and reduced, but not complete loss of all activity. Incomplete ablation may be due to insufficient Mtz (although we used the highest dose possible at this stage without inducing morphological defects). Another possibility is that mural cells are undergoing rapid differentiation at this time and there may be a pool of mural cells that have myogenic activity, but do not express enough of the marker to undergo ablation. A third possibility is that there is mosaic expression of the transgenes and not every mural cell is labelled (and therefore not targeted by Mtz ablation). Although there was a large overlap between our pericyte transgenics, there were a number of cells that were not marked by both lines. Transgenic lines for vSMCs are also relatively mosaic at 4 dpf. Despite these limitations, our evidence strongly suggests that at 4 dpf, pericytes are actively regulating vessel diameter.

It is important to note that there are diverse responses of individual cells to vasoactive agents. Inconsistent responses from individual pericytes has also been observed in the mouse model. Only a population of pericytes contract in response to the vasocontrictor U46610 in a cranial window model, with half of the capillaries constricting and the other half dilating (Fernandez-Klett et al., 2010). Additionally, while some pericyte-covered regions respond to vasoactive agents, other regions may dilate in response to accommodate for the blood flow within the vascular network. It is also possible that not all pericytes have the capacity to respond to vasoactive agents. For instance only a subset of pericytes have been observed to express smooth muscle actin (αSMA) in the brain (Bandopadhyay et al., 2001), spinal cord (Roufail et al., 1995), retina (Nehls, 2004), and pancreatic islets (Almaça et al., 2018).

### Mechanism of pericyte contraction and relaxation

Intracellular calcium concentration dictates whether pericytes will contract or relax, at least in mice (Sakagami et al., 1999; Sakagami et al., 2001; Sugiyama et al., 2004). Pericytes can contract in response to endothelial derived vasoactive agents (Matsugi et al., 1997) but do not express the same contractile machinery as vSMCs. Pericytes lack the calcium-binding protein calponin, which is responsible for regulating contraction in vSMCs (Bandopadhyay et al., 2001). However, pericytes do contain microfilaments resembling actin and myosin containing fibers (Ho, 2004; Lebeux and Willemot, 1978). In cultured pericytes, activation of endothelin-1 (ET-1) receptors in pericytes increases calcium, resulting in the alignment of F-actin and intermediate filaments and contraction (Dehouck et al., 1997). We used noradrenaline and phenylephrine to assess the contractile properties of pericytes during development because catecholamines such as noradrenaline have been shown to modulate tone in cultured pericytes (Markhotina et al., 2007), and in cerebral slices (Hall et al., 2014; Peppiatt et al., 2006).

Our understanding of pericyte relaxation is limited. Similar to vSMCs, pericytes relax by activation of potassium (Burnette and White, 2006; Cao et al., 2006; Jackson, 2005; Li and Puro, 2001; Quignard et al., 2003; Von Beckerath et al., 2000; Wiederholt et al., 1995). Activation of potassium channels by vasodilators such as NO reduces the activity of L-type voltage operated calcium channels (VOCC) and calcium activated chloride channels (Cl_Ca_) in pericytes (Sakagami et al., 2001). The mechanism by which NO causes pericyte relaxation appears to be through prostaglandin I2 activity, a vasodilator (Burnette and White, 2006; Dodge et al., 1991). The EP4 receptor contributes to pericyte relaxation (Hall et al., 2014) in conjunction with NO, which is needed to stop the production of vasoconstricting agent 20-HETE (Hall et al., 2014). We show that blocking the EP4 receptor followed by exposure to the NO donor SNAP results in constriction and blocks dilation, highlighting the importance of the prostaglandin pathway in pericyte relaxation in vivo.

### A temporal shift in cell types regulating vascular tone

The relationship between neural activity and pericyte activity is not well understood. Although pericytes can locally modulate capillary diameters resulting in local changes in red blood cell (RBC) flow, this activity is unrelated to neurovascular coupling according to one study (Fernandez-Klett et al., 2010). In another study, whisker pad stimulation results in robust calcium transients in pericytes, corresponding to changes in vessel diameter and blood flow, and pericytes rather than vSMCs were shown to initiate this process (Hall et al., 2014). However a third study showed that stimulation of neuronal function via whisker pad stimulation results in robust calcium transients only in vSMCs, and subsequent vSMC-covered vessel dilation (Hill et al., 2015). Hill et al. found delayed dilation in vessels without smooth muscle coverage, specifically after peak dilation in vessels with SMC coverage. Thus, this study suggested the dilation in non-vSMC covered vessels is a passive response to the upstream increase in blood flow (Hill et al., 2015).

Understanding when neurovascular coupling develops in an organism is crucial to elucidating the role of cerebral vascular mural cells in regulating vascular tone. When does neurovascular coupling develop in fish? Ulrich et al. determined that up until 3 dpf, the development of neuronal structures is not effected by the presence or absence of vessels in the zebrafish hindbrain (Ulrich et al., 2011). Furthermore, while there is increased neuronal activity at 6 dpf in the optic tectum in response to visual stimulus in fish larvae (Chhabria et al., 2018), it does not correlate to increased red blood cell speed (RBC) until 8 dpf, a time when embryos exhibit neurovascular coupling in the optic tectum. These time windows correlate with the two windows of mural cell activity in our data. At 4 dpf, while neurovascular activity and coupling is immature, pericytes are present in large numbers in cerebral vessels and regulate cerebral vessel diameter and hence, blood flow. This is further supported by our results that show physical contact between astrocytes and pericytes at 4 dpf, both components of the neurovascular unit. This suggests that at 6 dpf, pericytes are coordinated as part of the neurovascular unit. Subsequently, neurons and astrocytes could be inhibiting rather than stimulating pericyte activity in response to exogenous vasoactive agents. Once neuronal activity has increased, the coupling of neuronal activity and changes in blood flow matures.

Our findings in developing zebrafish embryos reveal key similarities and differences to data generated primarily in the adult mouse brain model. We show that both mural cell types are involved in regulating cerebral vessel diameters but at different stages of development. Pericytes, which share a developmental origin of neural crest with vSMCs in the forebrain and hindbrain (Ando et al., 2016; Cavanaugh et al., 2015; Wang et al., 2011; Wang et al., 2014; Whitesell et al., 2014), regulate cerebral vessel diameter and local changes in CBF while vSMCs are differentiating at 4 dpf. It is important to remember that although the vessels covered by pericytes are the smaller capillaries, RBCs have a diameter of ∼5 μm in the zebrafish and these cells have the ability to deform while squeezing through constricted blood vessel regions (Noguchi and Gompper, 2005; Pawlik et al., 1981; Skalak and Branemark, 1969). Thus, a small change in diameter on the smallest vessels might dramatically decrease resistance and increase blood flow.

Later, once vSMCs have matured and significant neuronal activity occurs, vSMCs become the primary regulators of vascular tone. Although, there are gap junctions between pericytes, endothelial cells and vSMCs (Borysova et al., 2013; Cuevas et al., 1984), there is no data in zebrafish to elucidate the mechanism by which neuronal activity causes changes in vessel diameter and blood flow. Do pericytes signal to upstream vSMC-covered vessels to respond to neural activity, or do molecular signals from neurons directly affect vSMC activity? These questions have yet to be answered. Clearly both pericytes and vSMCs are capable of responding to exogenous vasoactive agents, but the sources and developmental ontogeny of endogenous vasoactive signals remain unknown. Going forward, studies involving local activation of neural or mural cell activity will be needed to unravel the interactions between mural cell subtypes and developmental neurovascular coupling.

## Acknowledgements

This work was funded by a National Science and Engineering Council (NSERC) grant RGPIN/07176-2019 to SJC. NB is the recipient of a studentship from the Alberta Children’s Hospital Research Institute. We would like to thank Dr Jae-Ryeon Ryu, Dr. Thomas Richard Whitesell, Dr. Suchit Ahuja, Charlene Watterston, Jasper Greysson-Wong and Dr. Peng Huang and his laboratory for their insightful suggestions and helpful comments on this project. We would also like to thank Dr. Pina Colarusso and other members of the Live Cell Imaging Resource Lab at the University of Calgary for their microscopy help.

